# Two paralogues of N-ethylmaleimide sensitive factor: An exception to the minimal vesicular trafficking machinery of *Giardia*

**DOI:** 10.1101/2024.08.20.608744

**Authors:** Trisha Ghosh, Shankari Prasad Datta, Pritha Mandal, Nabanita Patra, Kuladip Jana, Sandipan Ganguly, Srimonti Sarkar

## Abstract

Vesicular trafficking plays a critical role in the survival of the human gut pathogen *Giardia lamblia* as it drives nutrient uptake and morphological stage transition. Unlike most eukaryotes, *Giardia* has a minimal vesicular trafficking machinery. Herein, we report a rare exception to this minimalism wherein two paralogues of NSF, a crucial factor driving vesicular trafficking by uncoupling the cis-SNARE bundle, are present in this unicellular parasite. While GlNSF_114776_ and GlNSF_112681_ share very high sequence homology, they are likely to have distinct cellular roles as they exhibit differences in their affinities towards the Glα-SNAPs and display non-overlapping distribution in encysting trophozoites. Under multiple stress conditions (nutritional, oxidative and nitrosative), while GlNSF_112681_ remains at peripheral vesicles, GlNSF_114776_ relocalizes to the anterior flagella-associated striated fibres, indicating a possible role in regulating flagellar motility. At this location, GlNSF_114776_ is likely to perform a 20S complex independent function as neither Glα-SNAPs nor GlSNAREs are present there. The two paralogues are likely needed for stress adaptation as both copies have also been retained in *Giardia* genomes isolated from clinical samples. This non-canonical function of the GlNSF_114776_ may have evolved to support the unique architecture and motility of the anterior flagella.

## Introduction

*Giardia lamblia*, a unicellular protist, is the most common cause of parasitic diarrhoea worldwide [1]. It has a two-stage life cycle: (i) trophozoites- a binucleated and flagellated form that colonizes the intestinal epithelium of the host and is responsible for the manifestation of the disease; (ii) cysts- a non-motile, tetra-nucleated, infectious form that transmits via the faecal-oral route [2]. The subcellular architecture of *Giardia* varies considerably from that of most eukaryotes [1]. The endomembrane system of this protist is relatively rudimentary [3]. Most eukaryotes have an endomembrane system comprising various compartments, *viz.*, the endoplasmic reticulum, Golgi, endosomes, and lysosomes. Transport vesicles mediate the exchange of various macromolecules between these compartments as they bud off from the donor compartment and fuse with the acceptor [4]. Based on electron micrographs, *Giardia*’s endomembrane system appears simpler, with only two types of compartments: the endoplasmic reticulum and the peripheral vesicles (PVs) [5]. PVs are the endo-lysosomal equivalent in this parasite [6]. They are distributed close to the plasma membrane and serve as entry sites through which the parasite uptakes nutrients from the host gut milieu [7]. Besides nutrient uptake, the endomembrane system is required for transitioning between the trophozoites and the cysts [8]. This interconversion between the two morphological states of *Giardia* is vital for the parasite’s survival inside the host gut and for its conversion to a form that can persist in the environment [9]. It is intriguing that even this minimal endomembrane system of *Giardia* can still fulfil the same functional roles that a much more complex endomembrane system discharges in most other eukaryotic organisms. Therefore, understanding the components of the endomembrane system of *Giardia* is crucial in elucidating how this minimalist system supports the parasite’s survival strategies.

As previously mentioned, vesicles ferry macromolecules from one compartment to another. Fusion between a vesicle and its target membrane necessitates that their lipid bilayers come very close to each other to allow the mixing of lipids [10]. The SNAREs (<u>s</u>oluble <u>N</u>SF [N-ethylmaleimide-sensitive factor] <u>a</u>ttachment protein <u>re</u>ceptors) are instrumental in enabling this close apposition of the two membranes [11]. Members of the SNARE family are relatively small proteins that contain a coiled-coil domain [12,13]. They are compartment-specific, with R-SNAREs on vesicles pairing with the cognate Q-SNAREs of the target membrane [14]. Selective distribution of various Q- and R-SNAREs to specific membranes and the highly selective pairing between them contributes to the specificity in the vesicle targeting process [15,16]. The strong interaction between the coiled-coil domains of the SNAREs within the trans-SNARE complex brings the two membranes very close, thereby driving membrane fusion [17]. Following fusion, the SNARE complex is termed a cis-SNARE complex, since it is now located on the same membrane. This hyper-stable complex must be disassembled so that the R-SNAREs can recycle back to their original compartment; this allows the same SNARE molecule to participate in multiple rounds of vesicle fusion.

The disassembly of the SNARE complex is accomplished by the action of a AAA ATPase, NSF, and its adapter, α-<u>s</u>oluble <u>N</u>SF <u>a</u>ttachment <u>p</u>rotein (α-SNAP) [18]. α-SNAP facilitates SNARE recycling by interacting directly with the cis-SNARE and the ATP-bound NSF [19,20]. Since NSF has no SNARE binding surface, α-SNAP mediates the interaction between NSF and SNAREs and stimulates the ATPase activity of NSF, ultimately leading to SNARE complex disassembly [21]. The complex comprising NSF, α-SNAP, and SNAREs is termed the 20S complex [22,23]. High-resolution cryo-electron microscopy of the 20S complex (resolution 7.35 Å) reveals that four copies of α-SNAP and six copies of NSF are present in a single 20S complex [24]. The α-SNAPs form a sheath around the SNARE bundle, upon which the NSF binds and uses energy released from ATP hydrolysis to disassemble the SNAREs [25–27]. Both NSF and α-SNAP are essential for sustaining eukaryotic vesicle fusion events as they make SNAREs available for further rounds of vesicle fusion.

As previously mentioned, there is limited endomembrane compartment diversity in *Giardia.* In addition, literature documents a reduction in the components of the machinery that sustains this rudimentary endomembrane system [28]. The *Giardia* genome encodes only two of the four adaptor protein complexes identified in model eukaryotes, and several components of the vesicle tethering complexes are absent [29,30]. Additionally, only eight Rab GTPases have been identified in *Giardia*, contrasting with the diverse array found in other parasitic protists such as *Trichomonas* and *Entamoeba* [31–35]. Compared to the larger repertoire of SNAREs present in most eukaryotes, *Giardia* has fewer SNAREs [36]. Moreover, the ESCRT machinery in *Giardia* consists of fewer components, with entire complexes like ESCRT-0 and ESCRT-I missing or those like ESCRT-II and ESCRT-III containing fewer subunits [37,38]. However, an exception to this minimalism has been previously observed in the case of α-SNAPs [39]. *Giardia* has three paralogues of α-SNAP, which are substantially diverged, not only from other eukaryotic α-SNAPs but also from each other (identities ranging between 21.6% to 30.6%). In addition to having multiple α-SNAPs, here we report the presence of two paralogues of NSFs in the reference genome of *Giardia* and also in clinical samples. The presence of such multiple paralogues of NSF is very rare in eukaryotic genomes. We have determined the subcellular localization of these individual NSF paralogues to understand if they play independent roles in this unicellular parasite. Our results indicate that while both paralogues will likely be involved in vesicle fusion events, at least one NSF may discharge a parasite-specific role during stress conditions.

## Results

### *Giardia* genome encodes two paralogs of NSFs

Searches of the *Giardia* genome indicated the presence of two putative NSF genes, encoded by ORFs GL50803_112681 and GL50803_114776, respectively. The presence of such paralogous genes encoding NSF is unusual, as most organisms have only one copy. We have previously reported the localization of one of these paralogues, GlNSF_114776_, in trophozoites and encysting trophozoites [39]. Sequence comparison of the two GlNSF paralogues indicated they share significant identity at the nucleotide (92.7%) and protein (89.7%) levels. This high degree of sequence identity makes it difficult to differentiate between the expression of these two genes. A perusal of the transcriptomics data sets available in GiardiaDB highlights this problem, as many of these results are annotated with a caveat regarding non-unique aligned reads. To circumvent this problem, we designed separate primer sets that would specifically amplify each GlNSF gene (Supplementary Figure 1 and Supplementary Table 1). We used these primers to perform reverse transcription PCR, using RNA isolated from trophozoites, encysting trophozoites and cysts. The results indicated the amplification of specific products whose sizes match the length of predicted amplicons (Figure 1A). This indicated that both genes are expressed in all the lifecycle stages included in this analysis.

**Figure 1:**
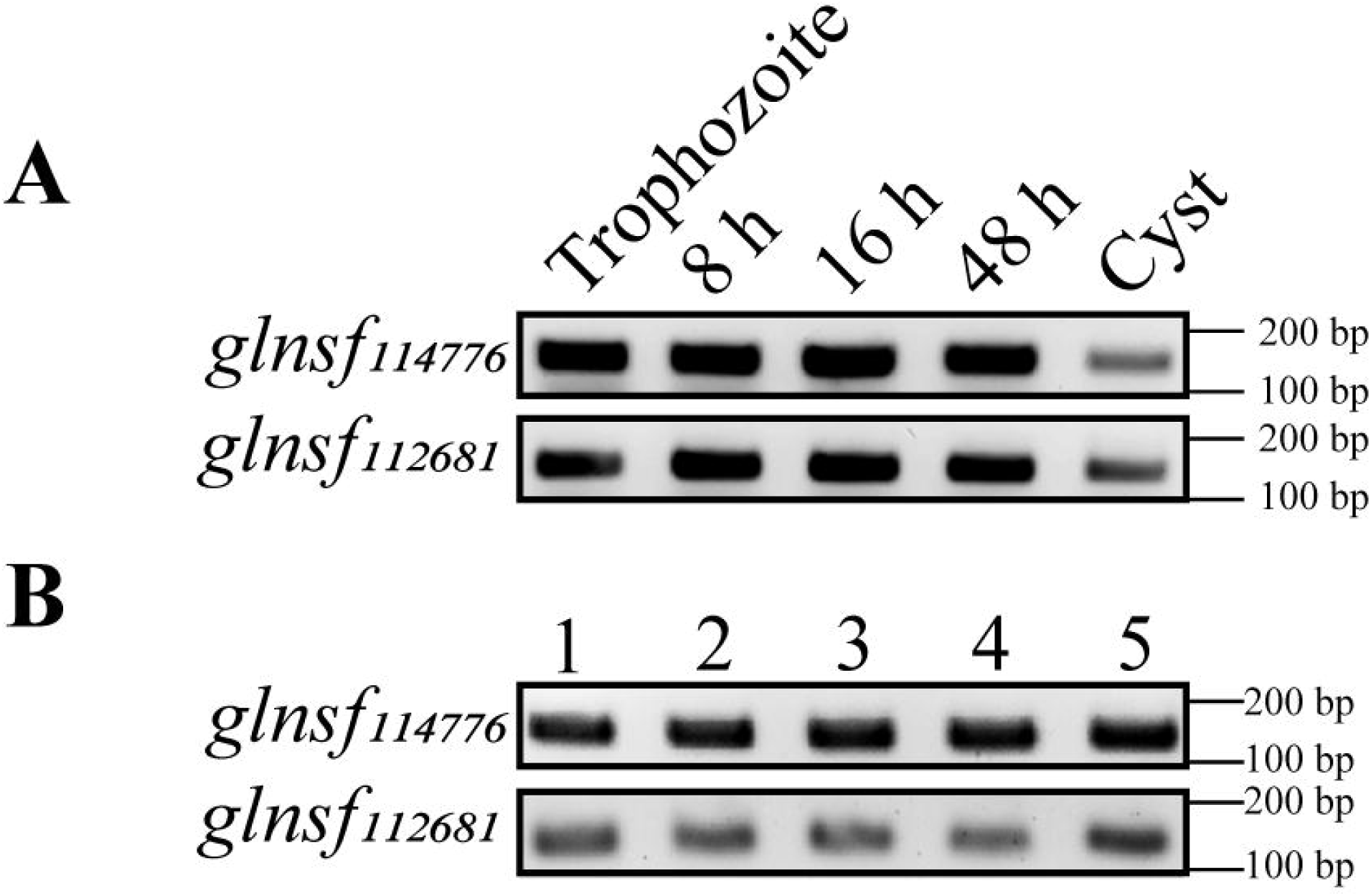
Expression of the two putative *glnsf* genes. (A) Expression of *glnsf_114776_* and *glnsf_112681_* in *Giardia* trophozoites, encysting trophozoites (8 h, 16 h, and 48 h) and cysts. (B) PCR amplification of *glnsf_114776_* and *glnsf_112681_*from clinical samples. PCR was performed with genomic DNA extracted from five different samples as template and gene-specific primers.

The presence of two paralogs of GlNSF was confirmed by mass spectrometry. Whole-cell extracts from *Giardia* trophozoites were subjected to LC-MS/MS analyses, and the peptides mapping to either paralogous sequence were identified. We considered only those peptide fragments that were detected at least thrice (Supplementary Tables 2 and 3). Some of these peptides mapped to regions common for both paralogues. However, many fragments could be specifically assigned to a given paralogue based on the presence of amino acid residues that are unique to each sequence. Sixty-six such fragments were mapped onto GlNSF_114776_, while sixty-one were mapped onto GlNSF_112681_. This mapping of unique sequences to each paralogue resulted in 60.43% sequence coverage for GlNSF_114776_ and 58.55% for GlNSF_112681_. Based on these results, we conclude that not only are the two paralogous genes encoding GlNSF transcribed, but they are also translated in trophozoites.

Besides establishing the presence of two paralogues of GlNSF in the reference strain of *Giardia*, we also determined if *Giardia* genomes isolated from clinical samples contain multiple paralogues. Genomic DNA was isolated from five clinical samples and used as templates for PCR amplification using GlNSF paralogue-specific primers. In all five cases, we observed specific PCR amplification of both loci (Figure 1B). Thus, based on the expression of both paralogues of NSF in different stages of *Giardia*’s life cycle, the detection of each individual protein in trophozoite extracts, and the presence of the two paralogous genes in multiple clinical samples, we conclude that both the paralogous proteins play a role in the biology of this human pathogen, and are likely to discharge specific cellular functions.

### GlNSF_112681_ resides at vesicles and colocalizes with all the Gl**α**-SNAPs

We have previously reported that GlNSF_114776_ is present at the PVs in trophozoites, where all three Glα-SNAPs localize either throughout or during certain stages of the parasite lifecycle [39]. This indicates that GlNSF_114776_ will likely discharge a Glα-SNAP-dependent function at the PVs. However, during the course of encystation, GlNSF_114776_ progressively migrates to the region of the cell occupied by striated fibres, which are repetitive structures associated with the cytoplasmic portions of the anterior flagella axoneme [39]. The functions of these striated fibres are not known [40]. Since the PVs are active sites of vesicular trafficking [5], and we have previously observed a significant reduction in the pool of GlNSF_114776_ at this site during encystation, we hypothesize that the paralogous GlNSF_112681_ will be present at PVs during encystation to sustain membrane fusion events at this location. To test this hypothesis, we determined the subcellular distribution of this GlNSF paralogue via immunolocalization.

The high sequence identity between the two GlNSF paralogues presented a challenge towards raising antibodies specific to each GlNSF. In our previous study, we had raised polyclonal antibodies against GlNSF_114776_ in rats [39]. To ensure this antibody does not bind to GlNSF_112681_, we determined its ability to differentiate between the two paralogous proteins expressed in *E. coli*. Both proteins were expressed with a 6xHis tag and purified. To rule out the possibility that the antibody recognizes the 6xHis tag, the tag was removed from GlNSF_114776_, but not GlNSF_112681_. The antibody specifically detected only the GlNSF_114776_, but not the 6xHis-tagged GlNSF_112681_, which proves that this polyclonal antibody is specific to GlNSF_114776_ and does not exhibit any affinity for either GlNSF_112681_ or the 6xHis tag (Figure 2A). Polyclonal antibodies specific for GlNSF_112681_ were raised against the N-terminal of this protein as multiple sequence alignment revealed the presence of unique regions within the N-terminal domains of these two paralogues (Supplementary Figure 2). The raised antibody detected a band close to 55 kDa in *E. coli* extract, consistent with the expected size of the tagged N-terminal fragment (Figure 2B). This antibody was specific to GlNSF_112681_ as it failed to detect bacterially expressed GlNSF_114776_. When the antibody was used for western blotting of *G. lamblia* trophozoite extract, only a single band of ∼91 kDa was observed, consistent with the predicted size of GlNSF_112681_ (Figure 2C). No signal was detected with the preimmune antisera. Taken together, these results indicate that even though the two GlNSF paralogues share considerable sequence identity, each of the two antibodies can specifically detect the target GlNSF. Hence, these antibodies were used for immunolocalization experiments to determine the subcellular distribution of each GlNSF.

**Figure 2:**
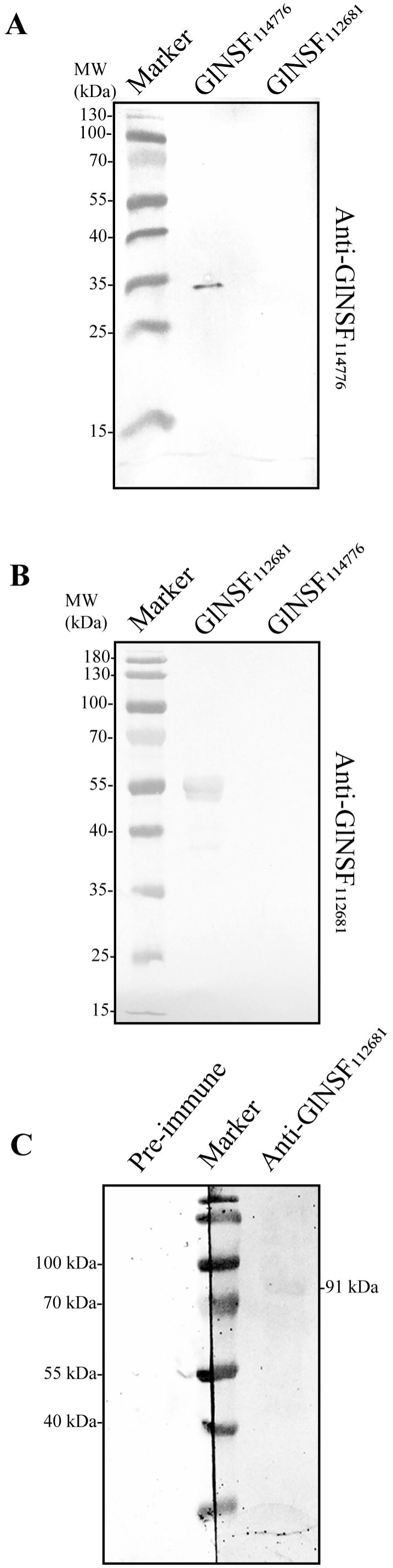
Western blotting to determine the specificities of antisera against the GlNSF paralogues. (A) The N domain of both GlNSF_114776_ and GlNSF_112681_ were expressed in *E. coli* with 6xHis-tag. The tag was removed from GlNSF_114776_ but not GlNSF_112681_ and probed with anti-GlNSF_114776_. (B) The same protein fragments bearing the 6x-His tag as in (A) were probed with anti-GlNSF_112681_. (C) Protein extract, prepared from *Giardia* trophozoites, was probed with either the pre-immune sera or anti-GlNSF_112681_.

Immunolocalization of GlNSF_112681_ in *Giardia* trophozoites indicated that it is primarily present at the PVs, which is similar to that reported for GlNSF_114776_ (Figure 3A) [39]. A few cytoplasmic puncta were also observed. There was a progressive increase in these cytoplasmic puncta during the 8-16 h period post-induction of encystation and a decrease in the number of PVs positive for this GlNSF (Figure 3A). Based on the small sizes of the observed cytoplasmic puncta, they do not appear to be encystation-specific vesicles [5]. At 48 h after induction of encystation, the signal at the PVs was similar to that at 16 h, but there was a reduction in cytoplasmic puncta at the later time point (compare Figure 3A and 3B). Although there is a decrease in the peripheral distribution of GlNSF_112681_ during encystation, this depletion is not as significant as that previously observed for GlNSF_114776_ [39]. Thus, consistent with our hypothesis, the presence of GlNSF_112681_ at the PVs will likely sustain the membrane fusion events at these compartments and compensate for the depletion of GlNSF_114776_ at this location.

**Figure 3:**
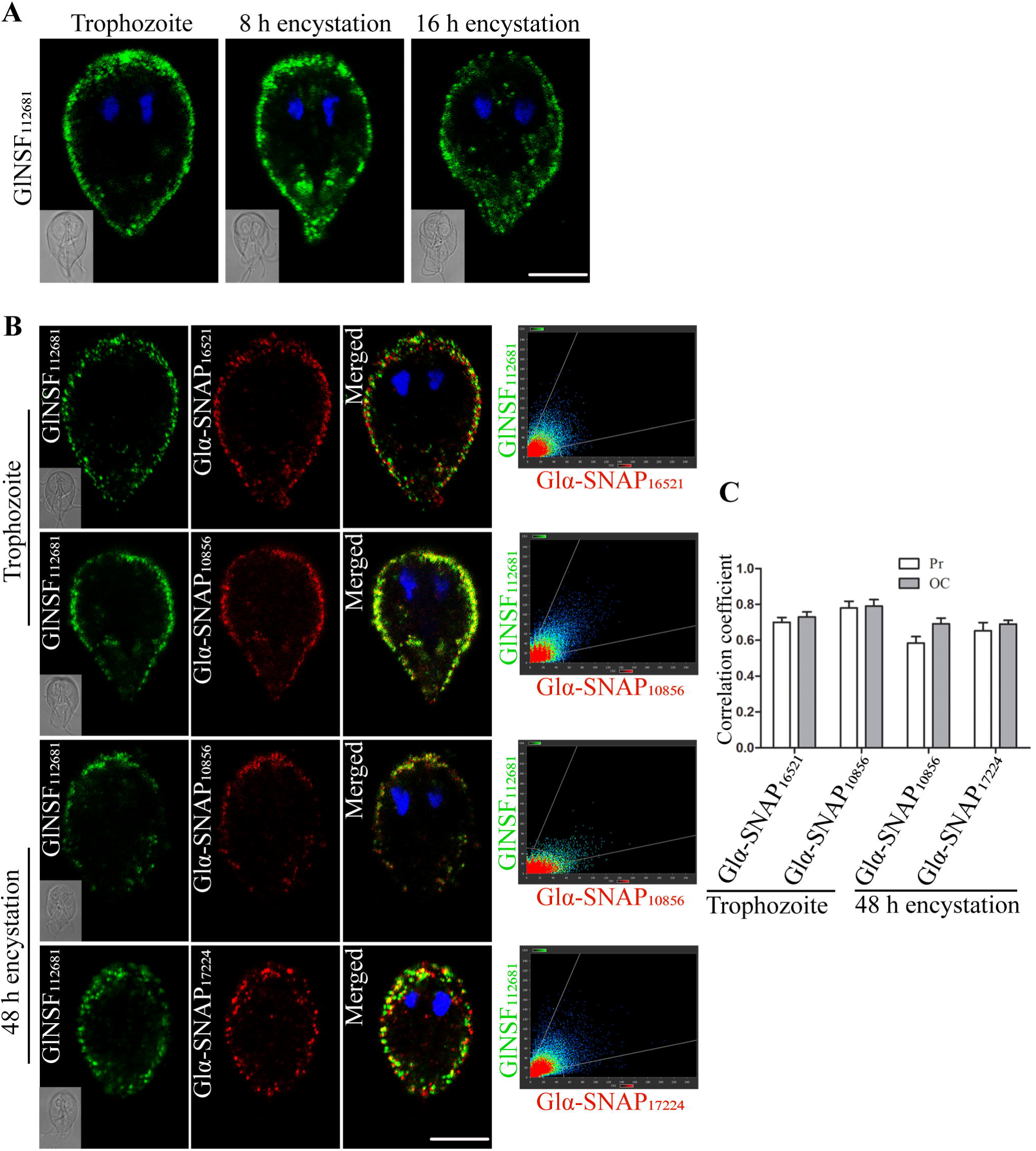
Subcellular distribution of GlNSF_112681_ and its colocalization with the paralogues of Glα-SNAPs. (A) Immunofluorescent detection of GlNSF_112681_ in trophozoites and encysting trophozoites (8 h and 16 h). DIC images are in the inserts. (B) Co-localization of GlNSF_112681_ with Glα-SNAP_16521_ or Glα-SNAP_10856_ in trophozoites and with Glα-SNAP_10856_ or Glα-SNAP_17224_ in 48 h encysting trophozoites. The colocalization analysis of the two fluorophores throughout the entire cell is depicted in each adjacent scattergram. (C) Mean Pearson’s correlation coefficient (Pr) and overlap coefficient (OC) values from the z-stacks of five independent images to determine the extent of colocalization of GlNSF_112681_ with Glα-SNAP_16521_ or Glα-SNAP_10856_ in trophozoites and after 48 h of induction of encystation with Glα-SNAP_10856_ or Glα-SNAP_17224_. *Scale bar* A, B 10 µm.

Since the formation of a functional 20S complex requires both α-SNAP and NSF, it is expected that a true GlNSF will colocalize with the Glα-SNAPs. We investigated whether the three paralogous Glα-SNAPs colocalize with GlNSF_112681_. As previously reported, all the Glα-SNAPs display a peripheral distribution during a certain stage of the parasitic life cycle [39]. While Glα-SNAP_10856_ is present at the PVs in trophozoites and during encystation, Glα-SNAP_17224_ is located at the perinuclear region in trophozoites and relocalizes to the PVs at 48 h post-induction of encystation. Glα-SNAP_16521_ localizes to PVs in trophozoites and migrates to the paraflagellar dense rods early in encystation. Since the PV association of GlNSF_112681_ remains unchanged, we investigated its co-localization with the various Glα-SNAPs under conditions when each of these three was present at the PVs. Specifically, we determined colocalization of GlNSF_112681_ and Glα-SNAP_17224_ 48 h post-induction of encystation, and with Glα-SNAP_16521_ and Glα-SNAP_10856_ in trophozoites. Since Glα-SNAP_10856_ remains at the PVs during encystation, we also studied its colocalization with GlNSF_112681_ in 48 h encysting cells.

Colocalization with Glα-SNAP_16521_ in trophozoites showed that while the signals for the two proteins overlapped at the cell periphery of trophozoites, this Glα-SNAP was also present in cytoplasmic puncta where the GlNSF_112681_ was absent (Figure 3B). Also, the two signals did not overlap significantly at the anterior edge of the trophozoite. However, at these locations, Glα-SNAP_16521_ may still function in conjunction with GlNSF, as we have previously observed that it has significant colocalization with GlNSF_114776_ in punctate structures in the cytoplasm and towards the anterior edge of the cell [39]. To quantify the extent of colocalization, we carried out quantitative image analysis and determined the values of Pearson’s correlation coefficient (Pr) and overlap coefficient (OC) for the entire volume of five individual cells, and the mean values are shown (Figure 3C). The Pr value of 0.7, and the OC of 0.73 indicate partial colocalization and are concordant with the representative image.

When GlNSF_112681_ was colocalized with Glα-SNAP_10856_ in trophozoites, the signals for the two proteins overlapped more than that for Glα-SNAP_16521_, both at the cell periphery and at cytoplasmic puncta (Figure 3B). This is also supported by quantitative image analysis where Pr and OC values were 0.78 and 0.79, respectively. Colocalization with Glα-SNAP_10856_ in 48 h encysting cells indicated a decrease in the overlap of the two signals as the Pr value was 0.58 and OC was 0.69. But, like Glα-SNAP_16521_, this Glα-SNAP may also function within the context of the 20S complex as we have previously observed significant peripheral colocalization between it and the pool of GlNSF_114776_ that persists at the PVs [39].

Overlap of the signal of GlNSF_112681_ with Glα-SNAP_17224_ at 48 h also indicated partial colocalization with Pr of 0.65 and OC of 0.69. Based on the above observations, we conclude that at the cell periphery, GlNSF_112681_ partially colocalizes with all the Glα-SNAPs as the mean values of both Pr and OC were higher than the cut-off of 0.5 (Figure 3C). Scattergram analyses of individual representative images also support this conclusion. Taken together, we conclude that GlNSF_112681_ is likely to be responsible for most of the SNARE uncoupling taking place at the cell periphery as there is significant colocalization between it and the various paralogues of Glα-SNAPs at this location.

### N-terminal sequence divergence of GlNSF paralogues contributes to a selective affinity for Gl**α**-SNAPs

Since all the paralogues of Glα-SNAP colocalized with GlNSF_112681_, we wanted to determine if these proteins can physically interact with each other. The yeast-two hybrid (Y2H) assay has been previously used to identify interactors of NSF [41,42]. Using this assay, we have previously reported that GlNSF_114776_ interacts strongly with Glα-SNAP_10856_ and weakly with Glα-SNAP_17224_ [39]. This GlNSF paralogue was included in the present assay as a positive control. The two GlNSFs were fused with the activation domain (AD) of the yeast transcription factor Gal4, and the three Glα-SNAPs were expressed as fusions of the DNA binding domain (BD) of the same transcription factor. Positive interactions between the NSF and α-SNAP orthologues of *Giardia* were identified by monitoring the growth of the transformants on histidine-leucine-tryptophan triple dropout medium containing 2.5 mM 3-amino-1,2,4-triazole (LTH 3-AT) and leucine-tryptophan-adenine triple dropout medium (LTA) as these monitor the expression of the *HIS3* and *ADE2* reporter genes, respectively. The results of the Y2H assay indicate that unlike GlNSF_114776_, which is known to interact with two Glα-SNAPs, GlNSF_112681_did not interact with any of the three Glα-SNAP paralogues as there was no growth on both LTH-3AT and LTA plates (Figure 4A). The observed variation in the interaction of the two GlNSFs with that of the Glα-SNAPs highlights the likelihood that the GlNSFs may have undergone functional divergence within this parasite. Although there appears to be no binary interaction between the three Glα-SNAPs and GlNSF_112681_, we cannot exclude interaction happening in vivo, where either additional cellular factors or post-translational modifications (PTMs) might play a role in stabilizing such interactions.

**Figure 4:**
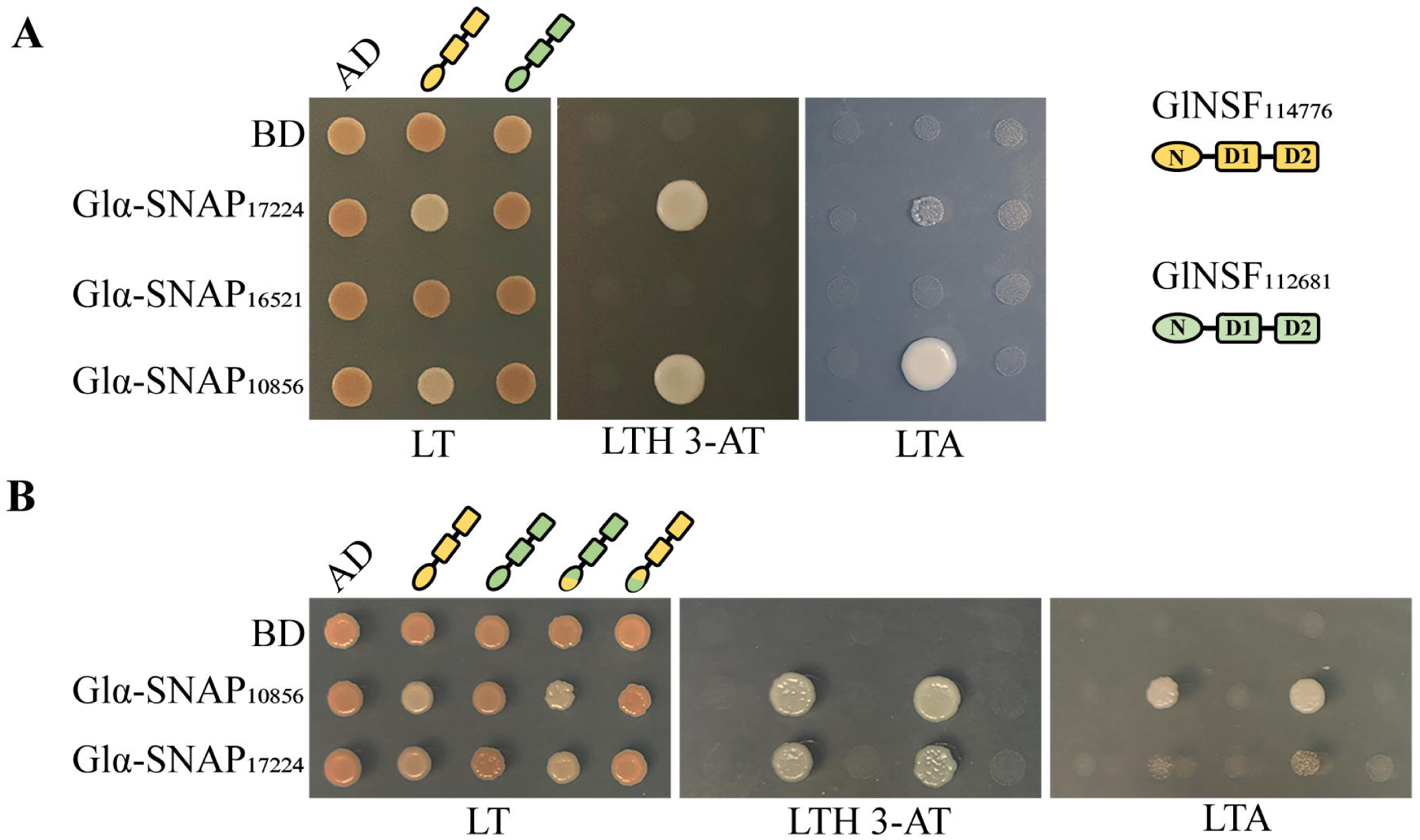
Binary interaction between GlNSF_112681_ and the paralogues of Glα-SNAPs. (A) PJ69-4A cells were transformed with various combinations of constructs expressing GlNSF_112681_ and GlNSF_114776_ as AD fusion proteins, while the Glα-SNAPs were expressed as BD fusion proteins. The transformants were spotted on YCM plates lacking leucine and tryptophan (LT), or leucine, tryptophan and histidine plates supplemented with 2.5 mM 3-amino triazole (LTH-3AT), or leucine, tryptophan and adenine (LTA). (B) PJ69-4A cells were transformed with varying combinations of constructs expressing the recombinant GlNSFs as AD fusion proteins and Glα-SNAPs as BD fusion proteins. The growth of these transformants were monitored on LT, LTH-3AT, and LTA plates. For both cases (A) and (B), the empty vectors AD and BD served as a negative control, while the transformant expressing GlNSF_114776_ and Glα-SNAP_10856_ served as a positive control.

To understand the basis of the selective interaction of only GlNSF_114776_ with the two Glα-SNAPs, we wanted to determine the region that contributes to this selective interaction even though the two paralogous proteins share a high degree of sequence similarity. Cryo-electron microscopy structure of the 20S complex shows that the N-terminal domain of NSF interacts with the C-terminal domain of α-SNAP [24]. We hypothesized that differences in the N-terminal region of the two paralogues contribute to this selectivity in interaction with the Glα-SNAPs. To test this hypothesis, we generated constructs that express recombinant GlNSFs with interchanged N-termini. The first ninety-five residues of GlNSF_114776_ were swapped with the corresponding one hundred and two residues of GlNSF_112681_; the difference in size arises because GlNSF_112681_ has a seven amino acid insertion within this region. These recombinant GlNSFs were used in Y2H assays to determine the changes in their interactions, if any, with Glα-SNAP_10856_ and Glα-SNAP_17224_ as only these two interact with GlNSF_114776_. We observed that the recombinant GlNSF with the N-terminal ninety-five residues of GlNSF_114776_ interacted with the two Glα-SNAPs even though the rest of this protein’s sequence was from GlNSF_112681_ (Figure 4B). Conversely, even though a major portion of the sequence of the other recombinant was from GlNSF_114776_, it did not interact with the Glα-SNAPs (Figure 4B). Thus, the selectivity in interaction between the NSFs and α-SNAPs of *Giardia* is dictated by differences in the N-terminal segment of the two GlNSF paralogues.

### The two GlNSF paralogues colocalize in trophozoites but not under stress conditions

Since both GlNSF_112681_ and GlNSF_114776_ are present at the PVs in trophozoites, we wanted to determine if they colocalize. Our results show a significant overlap in the distribution of these two proteins at the PVs of trophozoites (Figure 5A). In addition, both GlNSF_114776_ and GlNSF_112681_ were also present in cytoplasmic puncta in trophozoites. Scattergrams generated from multiple colocalization experiments showed significant overlap between the green and red pixels and the Pr and OC values were both 0.84 (Figure 5B). While these values are consistent with the images showing significant colocalization of the two proteins at the PVs, where major pools of both are present, the minor pools present in cytoplasmic puncta do not overlap. Based on this observation, we conclude that the two proteins may function together under certain conditions but have the ability to discharge independent cellular functions.

**Figure 5:**
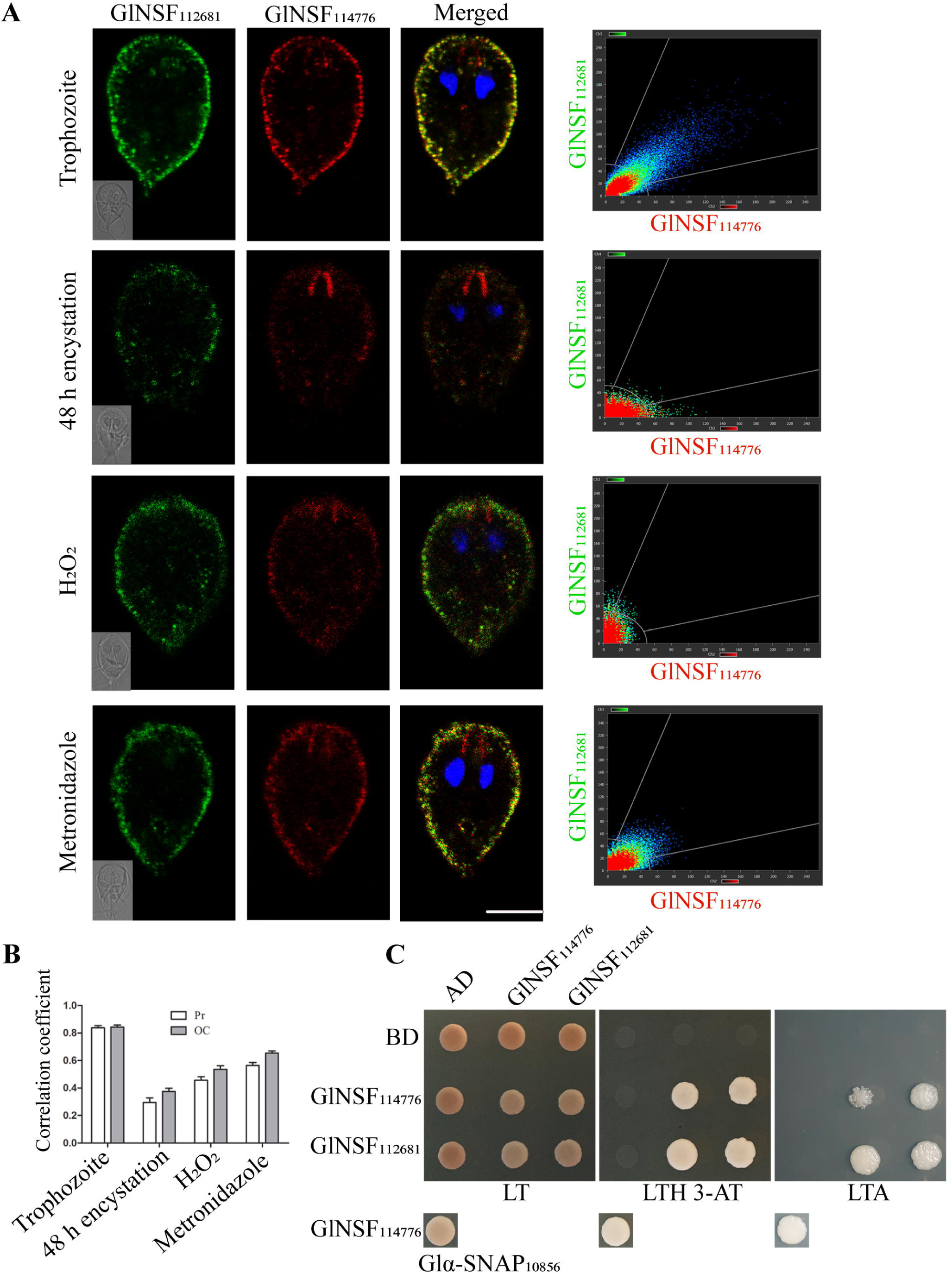
Co-localization and binary interaction between the paralogues of GlNSFs. (A) Colocalization of GlNSF_112681_ and GlNSF_114776_ in trophozoites, 48 h encysting trophozoites, and in trophozoites subjected to either oxidative or nitrosative stress. Oxidative and nitrosative stress was induced by treatment with 150 µM H_2_O_2_ or 1µg/ml metronidazole for 1 h. DIC images are in the inserts. The colocalization analysis of the two fluorophores throughout the entire cell is depicted in each adjacent scattergram. (B) The mean Pr and OC values calculated from the z-stacks of five individual cells are presented in the bar graph. (C) Binary interaction between the two GlNSF paralogous using a yeast-two hybrid analysis. PJ69-4A cells were transformed with different combinations of constructs expressing GlNSF_112681_ and GlNSF_114776_ as both AD and BD fusion proteins. The growth of these transformants was monitored on LT, LTH-3AT and LTA plates. The transformant harbouring the empty AD and BD vector was the negative control. The interaction GlNSF_114776_ and Glα-SNAP_10856_ served as positive control. *Scale-bar*, A 10 µm.

We have previously reported the GlNSF_114776_ migrates to the striated fibres of the anterior flagella during late encystation [39]. To determine if GlNSF_112681_ is capable of similar relocalization, we carried out colocalization of the two GlNSF paralogues 48 h after the induction of encystation and observed that while a major pool of GlNSF_114776_ migrated from the PVs to the striated fibres, GlNSF_112681_ remained at the PVs (Figure 5A). Images of 48 h encysting cells revealed that, in contrast to trophozoites, there was minimal overlap of the two fluorophores in these cells. Consistently, quantitative image analysis revealed low values of both Pr and OC, 0.30 and 0.38, respectively (Figure 5B). It may be noted that while a minor pool of GlNSF_114776_ remained at the PVs, there was no overlap between it and GlNSF_112681_; this is in contrast to the almost complete overlap observed in trophozoites (Figure 5A). Thus, while the two paralogues colocalize at the PVs of trophozoites, they exhibit distinct, non-overlapping subcellular distribution in late encysting cells.

As mentioned, a major pool of GlNSF_114776_ relocates from PVs to the striated fibres during encystation [39]. However, its subcellular distribution under stress was not assessed. Hence, we wanted to determine if both the GlNSF paralogues undergo relocalization under stress conditions. We carried out colocalization studies in trophozoites that were exposed to either the commonly-used anti-giardial drug, metronidazole, which results in both oxidative and nitrosative stress, or H_2_O_2_, which is a known oxidative stress-inducing agent [43–45]. Since *Giardia* lacks several crucial components employed by other organisms to detoxify reactive oxygen and nitrogen species, it is sensitive to H_2_O_2_ and metronidazole [46,47]. We determined the subcellular distribution of the two proteins after 1 h treatment with either 150 μM H_2_O_2_ or 1 μg/ml metronidazole [39,48]. Exposure to each reagent triggered intracellular oxidative stress, as evidenced by the conversion of DCFDA to the fluorescent DCF (Supplementary Figure 3). H_2_O_2_ exposure resulted in the relocalization of a pool of GlNSF_114776_ from the PVs to the striated fibres, while GlNSF_112681_ remained at the PVs (Figure 5A). Consequently, there is a significant reduction in the colocalization between the two paralogues, with the Pr and OC values being 0.46 and 0.54, respectively (Figure 5B). The treatment with metronidazole also caused similar relocalization of only GlNSF_114776_ (Figure 5A). Scattergram analysis revealed that, like H_2_O_2_ stress, exposure to metronidazole significantly reduced the colocalization of the two proteins, with Pr and OC values of 0.56 and 0.65, respectively (Figure 5B). These Pr and OC values are considerably less than the Pr and OC of the two GlNSF paralogues in unstressed trophozoites, which were both 0.84 (Figure 5A). Thus, only GlNSF_114776_ displays subcellular redistribution in response to oxidative stress, nitrosative stress, and encystation. Although the paralogous GlNSF_112681_ is very similar in sequence, there is no significant change in its cellular localization during the stress conditions or encystation. Based on these results, we conclude that even though the two GlNSF paralogues share a high degree of sequence identity, they are likely to perform non-overlapping functions during stress conditions.

The significant colocalization of the two GlNSF paralogues in trophozoites raises the possibility that the two proteins may be part of the same 20S complex, which has six copies of NSF [24]. To address this, we used a Y2H assay to determine if the two proteins could interact. The two GlNSFs were expressed in yeast fused with AD or BD at their N-termini. Transformants bearing all four different binary combinations of these fusion proteins were assayed for their ability to turn on the expression of the *HIS3* and *ADE2* reporter genes. All four transformants turned on the expression of the *HIS3* reporter as growth was observed on LTH 3-AT plates, where none of the negative controls survived (Figure 5C). Although the growth of the four different transformants was indistinguishable on LTH 3-AT plates, the growth pattern differed on LTA plates indicating varying affinities (Figure 5C). While both the AD and BD fusions of GlNSF_112681_ showed strong interaction with either the oppositely tagged GlNSF_114776_ or its own self, the interaction between the two GlNSF_114776_ was weaker. Based on the observed interactions between the two paralogous GlNSFs and their colocalization in trophozoites, it is likely that a heterogeneous 20S complex exists in *Giardia*.

## Discussion

We had previously reported the characterization of an NSF from *Giardia*. In the current study, we report that this unicellular eukaryote encodes another NSF paralogue. The data indicates that the newly identified GlNSF is also likely to function as a component of the 20S complex as it exhibits significant colocalization with the three Glα-SNAPs and is present at the PVs, where membrane fusion events occur frequently. Given the functional similarity between both the paralogues of GlNSFs, their retention in not only the reference strain but also in clinical samples indicates that they are likely to perform distinct and non-redundant roles during the lifecycle of this parasitic protist. This is particularly intriguing given that their sequences are very similar. The difference in the interaction of these two paralogous GlNSFs with the Glα-SNAPs, and their distinct localization under stress conditions further underscore their functional difference. This differential subcellular distribution indicates that the cellular machinery of this protist can distinguish between these two proteins and utilize them for parasite-specific functions.

Such a function seems likely for GlNSF_114776_ as we had previously observed that it undergoes subcellular relocalization during encystation and stress conditions, with a pool moving to the striated fibres associated with the anterior flagella and some being retained at the PVs [39]. Since none of the Glα-SNAPs or GlSNAREs is reported to be present at the striated fibres [36], GlNSF_114776_ likely discharges a non-canonical 20S complex-independent role at this particular location. However, the idea was tenuous since it raised concerns about how the PVs would function with only a minor pool of GlNSF. With the identification of a second GlNSF that remains at the PVs, there is additional support for a 20S-independent function of GlNSF_114776_. Such 20S independent roles of NSF have been previously documented, such as its direct interaction with the AMPA receptors for their recycling in the post-synaptic neuron and its binding to the C-terminal tail of the GluR2 receptors, which is required to maintain synaptic plasticity [41]. NSF can directly bind to the GluR2 independently of α-SNAPs, and all three domains of NSF are required for this activity [49,50]. The putative non-canonical function of GlNSF_114776_ at the striated fibres most likely arose to support the unusual biology of *Giardia*. These unique structures are associated with the anterior flagella axoneme, and other examples of such novel axoneme-associated structures are also observed for the other flagella [40]. These are thought to endow unique identities to each flagella pair, most likely enabling each pair to perform unique movements. While the morphology of the striated fibres has been revealed through SEM imaging, the factors contributing to their formation and sustenance remain unknown [40]. GlNSF_114776_ is the first protein reported to localize to this structure. Further characterization of its function at this location will elucidate how this unique cellular structure forms. This subcellular redistribution of GlNSF_114776_ occurred during encystation and when trophozoites were exposed to oxidative and nitrosative stress (Figure 5A). Such a response indicates that this protein may regulate flagellar activity under stress conditions.

PTMs may contribute to the rapid change in subcellular distribution of GlNSF_114776_ upon exposure to stress. Existing literature documents the role of PTMs in modulating the activity of the 20S components [51,52]. Phosphorylation and S-nitrosylation are known to modulate the interaction of NSF with the other components of the 20S complex. Phosphorylation of the Tyr83 and Ser237 inhibits the association of NSF with the SNAP/SNARE complex, while S-nitrosylation of its cysteine residues hinders the ATPase activity of NSF [53–55]. Our unpublished LC-MS/MS data also identified various PTMs in GlNSFs and Glα-SNAPs. We are currently investigating if some of these PTMs might modulate the interaction of the GlNSFs with the various paralogues of Glα-SNAPs. These PTMs may also uncover crosstalk between the vesicular trafficking system and other cellular processes.

Interestingly, while major pools of both GlNSF paralogues are observed at the PVs, there is a signal loss for both proteins during encystation (Figure 5A). This decrease indicates a possible reduction in membrane fusion events at the PVs, consistent with literature documenting their role as gateways of nutrient entry through bulk-phase fluid endocytosis [56]. As cells prepare to encyst, there is a switch in the metabolic circuitry of the cells wherein the focus shifts from nutrient uptake for growth and cell division to the formation of metabolically quiescent cysts. In comparison to the number of 20S complex-dependent membrane fusion events likely to be needed for nutrient uptake by trophozoites, that for the exocytosis of cyst wall material during encystation is likely to be fewer as the fusion of a single large encystation-specific vesicle with the plasma membrane will successfully deliver a considerable amount of cyst wall material to the cell surface [8].

We have previously reported that the three different α-SNAPs of *Giardia* colocalize with GlNSF_114776_.[39] In this study, we show that GlNSF_112681_ also colocalizes with the three Glα-SNAPs (Figure 3B). In addition, the two GlNSFs also exhibit significant colocalization at the PVs in trophozoites, but not in cytoplasmic puncta (Figure 5A). Based on the colocalizations of multiple components whose orthologues are known to be part of the 20S complex, we propose that the unusual morphology and unique subcellular architecture of this human parasitic protist may entail the formation of both location-specific pools of the 20S complexes, wherein different 20S complexes within the same cell are formed with different GlNSF paralogues, and heterogeneous 20S complexes composed of multiple GlNSFs and/or Glα-SNAPs. Investigating the 20S complex of this parasite in greater detail will shed light on the composition of this complex which in turn will give a better understanding of the unique vesicular trafficking machinery of this human parasite.

## Materials and Methods:-

### Sequence alignment of the GlNSFs

The protein and the cDNA sequences of both the GlNSF paralogues were curated from GiardiaDB. Sequence alignment was done using ClustalW, and visualization and editing were carried out using Jalview [57,58].

### Axenic culture of *G lamblia*

*G lamblia* (strain ATCC 50803/ WB clone C6) trophozoites were maintained in TYI-S-33 medium (pH 6.8) as previously described and encystation was induced as per established protocol [59,60].

### cDNA preparation and reverse transcription PCR

RNA was isolated from trophozoites, encysting trophozoites, and cysts, as described previously, and cDNA was prepared using reverse transcription PCR [61]. For selective amplification of the two GlNSF paralogous genes, primers specific to each paralogous gene were designed. The pair-wise nucleotide sequence alignment of the two paralogous genes revealed the presence of a unique 21-nucleotide segment in *glnsf*_112681_ which was used to design the forward primer; the reverse primer was designed to bind to a sequence stretch with four mismatches between the two paralogues (Supplementary Figure 1). The *glnsf*_114776_ sequence does not contain any unique stretch long enough to design a specific primer. However, there is a 13-nucleotide stretch near its 3’ end with six mismatches with the *glnsf*_112681_ sequence (Supplementary Figure 1). We designed an 18-mer reverse primer having four mismatches with this region of *glnsf*_114776_; these mismatches were not in any of the positions occupied by the previously mentioned six mismatches between the two paralogues. This primer was expected to bind to *glnsf_114776_* as there were only four mismatches in the 18-mer primer binding site, but it should not bind to *glnsf*_112681,_ as ten mismatches would not allow primer annealing. The forward primer was against a region present in both the paralogous genes (Supplementary Figure 1). Primer sequences are provided in Supplementary Table 1. After confirming the specificity of the primers, the cDNA was used to check the expression of both the paralogues during various stages of the parasitic lifecycle.

### Cell lysis and sample preparation for LC-ESI-MS/MS

A 250 ml confluent trophozoite culture was chilled on ice for 30 min and harvested by centrifugation at 1000g for 15 min. The cells were then washed twice with cold 1xPBS at 1000g for 10 min and resuspended in 200 µl lysis buffer containing 50 mM Tris pH 8.0, 120 mM NaCl, 5 mM EDTA, 1% Triton X-100, and a protease cocktail inhibitor (Sigma P8215). The cells were subjected to sonication (25 s pulses at 20% power followed by a 1 min rest period) [62]. This was repeated five times, and then the cells were incubated on ice for 40 min. The whole cell lysate was recovered by centrifugation at 13000 rpm for 10 min and lyophilized. This lyophilized protein was dissolved in 20 µl 100 mM NH_4_HCO_3_ (HiMedia). Bradford assay was performed to determine the lysate concentration and sample preparation for mass-spectrometry was done with 2.5 µg/µl protein. 25 µl trifluoroethanol (SRL) and 1.25 µl 200 mM DTT (Roche) were added to the sample and incubated for 1 h at 60°C. Subsequently, 5 µl 200 mM iodoacetamide (Sigma) was added, and the sample was incubated in the dark at room temperature (RT) for 90 min. Following this 1.25 µl 200 mM, DTT was again added to the sample and incubated for 1 h at RT. Finally, 219 µl water and 100 mM NH_4_HCO_3_ were added to the sample. The sample was digested overnight with 2.5 µl of 1 µg/µl trypsin (Promega V528A) at 37°C. The following day, the samples were centrifuged at 13000 rpm for 10 min, and the clear supernatant was collected. 1 µl 0.1% formic acid was added to the supernatant, and the sample was loaded for LC-ESI-MS/MS (Waters).

### LC-ESI-MS/MS data analysis

Three biological replicates were used to analyze the data. LC-MS/MS analysis was conducted using Progensis QI software for data acquisition and processing. The software recorded the mass spectra of the eluted peptides. Peptide identification was achieved through database searches against known protein sequence data from the UniProt database. Since both the GlNSFs share high sequence identity, they had many shared peptide fragments that could not be aligned to either of the protein sequences and hence, the software could not distinguish them as two separate proteins. Peptide fragment mapping onto both GlNSF protein sequences resolves this issue, identifying protein fragments with specific amino acid substitutions that make it unique to a specific GlNSF. Peptide fragments appearing at least thrice were considered during mapping, ensuring robustness. The protein sequence coverage for each GlNSF was calculated using only the uniquely aligned reads.

### Genomic DNA isolation and detection of GlNSF paralogues in clinical samples

Genomic DNA was extracted from patient stool samples using the QIAamp Fast DNA stool mini kit (Qiagen) following the manufacturer’s protocol. The presence of *Giardia* in clinical samples was determined as described previously by PCR using primers that targeted the β-giardin gene [62]. Once confirmed, the genomic DNA from these five different clinical samples was used as a template in PCR with gene-specific primers to verify the presence of the two GlNSF paralogues. All the experiments were conducted following the rules and regulations of the Institutional Human Ethics Committee of the Indian Council of Medical Research-National Institute of Cholera and Enteric Diseases (IRB Number: A-1/2015-IEC).

### Raising polyclonal antibodies against Glα-SNAP_10856_ and GlNSF_112681_

The genes for both Glα-SNAP_10856_ and GlNSF_112681_ were cloned in pET32a (Novagen) using the primers listed in Supplementary Table 1. Details of clone constructs are mentioned in Supplementary Table 4. Full-length Glα-SNAP_10856_ was expressed from BL21 (DE3) with 0.2 mM IPTG (GoldBio) at 30°C. The protein was purified from the soluble fraction as described previously to raise polyclonal antibodies in Wistar rats [38]. The N-terminal 312 amino acids of GlNSF_112681_ was expressed from Rosetta (DE3) with 0.1 mM IPTG at 37°C and was purified from the pellet fraction with 8M urea as described previously to raise polyclonal antibodies in Balb/C mice [63]. The animal experiments followed the protocols approved by the IAEC at Bose Institute, Kolkata (IAEC/BI/016/2022). Antibodies were purified from the collected sera using the NAb Protein A Plus Spin Kit (Thermo Scientific 89948) following the manufacturer’s protocol. Western blots were performed using trophozoite whole-cell lysate, as described previously, to assess the quality of the antibodies raised [61]. The pre-immune sera did not detect any bands in western blots (Supplementary Figure 4B). Moreover, when used in immunofluorescence experiments, the pre-immune sera did not show any non-specific localization for both cases (Supplementary Figure 4A and 4D). The polyclonal antibody against Glα-SNAP_10856_ displayed specificity without any cross-reactivity towards other Glα-SNAP paralogues (Supplementary Figure 4C).

### Immunofluorescence

Immunofluorescence experiments were performed on trophozoites and encysting trophozoites. Confluent cells were harvested, washed with 1xPBS, and fixed with 4% formaldehyde at RT for 15 min. Following this, the cells were washed with 1xPBS and treated with 1.5 M glycine for 10 min. Subsequently, the cells were washed with 1xPBS and permeabilized with 0.1% Triton X-100 for 10 min at RT, followed by blocking with 2% BSA in 1xPBS for 2 h at RT. Primary antibodies were used at 1:50 dilution in 0.2% BSA and incubated overnight at 4°C on a rocker at low speed. The next day, the cells were washed thrice with 1xPBS and incubated with different combinations of secondary antibodies as required at 1:400 dilution in 1xPBS for 1.5 h at RT on a rocker at low speed. The secondary antibodies used in this study were Alexa Fluor 594-conjugated goat anti-rat (Abcam-ab150160), Alexa Fluor 488-conjugated goat anti-mouse (Abcam-ab150113), and Alexa Fluor 594-conjugated goat anti-rabbit (Abcam-ab150080). Following this, the cells were incubated with 1µg/ml DAPI (Invitrogen D1306) for 30 min at RT and then washed thrice with 1XPBS. Subsequently, the cells were reconstituted in an antifade solution (0.1% p-phenylenediamine in 70% glycerol) and examined under a confocal laser scanning microscope (Leica, Stellaris) with a 63x objective. Images were processed using the Leica Application Suit X and all images were assembled using Adobe Photoshop CS3 and Adobe Illustrator CS3.

### Image analyses

Co-localization image analysis was done using the Leica Application Suit X. Correlation coefficients, including Pearson’s (Pr) and overlap (OC), were computed for each co-localization experiment by analyzing pixel-wise correlations between the signals emitted by two fluorophores across the z-stacks for five individual cells for each experiment. The mean Pr and OC values were represented in a bar graph using GraphPad Prism 5.

### Recombinant GlNSF constructs

*glnsf_112681_* was cloned in pGAD424 using the primers listed in Supplementary Table 1. Bsu36I, a restriction site common to both *glnsf* genes was used to create the recombinant *glnsf* constructs. Clones of the two *glnsf* paralogues in pGAD424 (cloned using EcoRI and SalI), were subjected to restriction digestion with EcoRI and Bsu36I. The gene fragments encoding the N-termini were swapped between the two *glnsf* paralogues. The clones were confirmed by sequencing. Details of the clone constructs are mentioned in Supplementary Table 4.

### Yeast two-hybrid assay

Glα-SNAPs and GlNSFs expressed as BD fusion and AD fusion proteins were used for the Y2H assay. The AD fusion protein was created by cloning the genes of interest in the prey vector pGAD424 (Clontech laboratories), having the *LEU2* selection marker, and bait vector pGBT9 (Clontech laboratories), having the *TRP1* selection marker, was used for creating the BD fusion protein [64,65]. Binary combinations of the various AD and BD constructs were co-transformed into the yeast strain PJ69-4A with *HIS3*, *ADE2*, and *lacZ* reporter genes under the control of three different galactose-inducible promoters from *GAL1, GAL2,* and *GAL7*, respectively. The transformants were selected on various yeast complete media (YCM) plates lacking leu-trp (LT-). To study the interactions among the various fusion proteins, the growth of PJ69-4A transformants on leu-trp-ade (LTA-) and leu-trp-his plates supplemented with 2.5 mM 3-amino triazole (LTH-3AT) was monitored after incubation at 28°C for 3 days. Cells transformed with the empty vectors served as a negative control. Details of clone constructs are mentioned in Supplementary Table 4.

### Induction of oxidative and nitrosative stress in *Giardia*

The growth medium of confluent trophozoite cells was replenished with fresh media and the cells were incubated at 37°C for 2 h. Oxidative and nitrosative stress was induced by treating the cells for 1 h at 37°C., with 150 μM H_2_O_2_ (Merck H1009) or 1 μg/ml metronidazole (Sigma M1547) respectively. To ascertain intracellular reactive oxygen species generation, the cells were chilled on ice and harvested by centrifugation at 1000g for 10 min followed by three 1xPBS washes. Subsequently, the cells were treated with 1.5 μM 2’,7’-dichlorodihydrofluoresceine diacetate (H_2_DCFDA) (Sigma D6883) for 15 min at 37°C. Following this, the cells were fixed with 2% paraformaldehyde, washed twice with 1xPBS, and then examined under a 63x objective of a confocal scanning laser microscope (Leica, Stellaris).

## Acknowledgement

This study was supported by funds from the Bose Institute and Anusandhan National Research Foundation (CRG/2022/001594/BHS). Fellowship support: TG from UGC (721/(CSIR-UGC NET DEC.2018)), SPD from INSPIRE (IF120153), PM from CSIR (09/015(0545)/2019-EMR-I) and NP from CSIR (09/015(0534)/2019-EMR-I).

## Data Availability Statement

This article and its supplementary files include all data generated or analysed during this study.

## Ethics approval and patient consent statement

All animal experiments were carried out under the approval of the Institutional Animal Ethics Committee, Bose Institute, Kolkata (IAEC/BI/016/2022). All experiments involving patient samples followed the regulations approved by the Institutional Human Ethics Committee of the Indian Council of Medical Research-National Institute of Cholera and Enteric Diseases (IRB Number: A-1/2015-IEC). Written consents were taken from all patients.

## Synopsis

*Giardia*’s minimal machinery for vesicular trafficking is well documented. Contrary to this minimalism, we document the presence of two NSF paralogues, a rare occurrence in eukaryotes. Even though these paralogues are highly homologous, their subcellular distributions indicate that they are likely to function independently, with one paralogue likely to participate in *Giardia*’s stress response. This study highlights the unique customizations of the vesicular trafficking machinery of *Giardia* that may have evolved to sustain the novel cellular features of this parasite.

## Abbreviations

PV: Peripheral vesicle
SNARE: Soluble NSF attachment protein receptor
NSF: N-ethylmaleimide sensitive factor
α-SNAP: α-Soluble NSF attachment protein
Pr: Pearson’s correlation coefficient
OC: overlap coefficient
AD: Activation domain
BD: Binding domain
PTM: Post-translational modification

## Acknowledgement

Confocal microscopy, DNA sequencing, and LC-ESI-MS/MS were conducted at the Central Instrument Facility of Bose Institute. We acknowledge the assistance of Prantik Saha, Sheolee Ghosh Chakraborty and Ayantrisha Biswas for confocal imaging, Mrinal Das for DNA sequencing, and Souvik Roy for mass spectrometry. We thank Prof. Alok K Sil for his suggestions and for proofreading the manuscript. We also thank all members of the Sarkar Laboratory for their valuable input. This study was supported by funds from the Bose Institute and Anusandhan National Research Foundation (CRG/2022/001594/BHS). Fellowship support: TG from UGC (721/(CSIR-UGC NET DEC.2018)), SPD from INSPIRE (IF120153), PM from CSIR (09/015(0545)/2019-EMR-I) and NP from CSIR (09/015(0534)/2019-EMR-I).

## Supplementary Figure Legends

Supplementary Figure 1: Pairwise sequence alignment of the two *glnsf* paralogues and location of the primers used for the selective amplification of each GlNSF paralogue. The red boxes denote the forward and reverse primer sequences for ORF GL50803_112681 designed from the mismatched region, while the black boxes denote the forward and reverse primer sequences for GL50803_114776 where point mutations have been incorporated in the reverse primer. The position of these point mutations has been underlined in the primer list provided in Supplementary Table 1.

Supplementary Figure 2: Primary structure alignment of the GlNSF paralogues. The asterisks denote positions of residues with change in the charge (positive ↔ negative or neutral ↔ charged) in the N-terminal domain of the GlNSFs. GlNSF_112681_ has an insertion of seven amino acids at the N-terminus.

Supplementary Figure 3: Intracellular reactive oxygen species generation, post-treatment with H_2_O_2_ or 1µg/ml metronidazole for 1 h, was assessed in trophozoites using DCFDA. Cells exposed to either H_2_O_2_ or metronidazole exhibited DCF fluorescence (green colour), indicating reactive oxygen species DCFDA generation, compared to the lack of green signal in untreated cells. *Scale-bar* 10 µm.

Supplementary Figure 4: Specificity of raised polyclonal antibodies. (A) Immunofluorescence studies using the pre-immune sera from mice and anti-GlNSF_112681_. (B) Anti-Glα-SNAP_10856_ raised in rats detected a single band of ∼33 kDa in trophozoite whole-cell extracts. No band was detected with the pre-immune sera. (C) Anti-Glα-SNAP_10856_ detects Glα-SNAP_10856_ alone, but none of the other two Glα-SNAPs, in *E. coli* extracts expressing each individual Glα-SNAP. (D) Immunofluorescence studies using the pre-immune sera from rats and anti-Glα-SNAP_10856_. *Scale-bar* A, D 10 µm.

