## Supplementary figures and images for "Two paralogues of N-ethylmaleimide sensitive factor: An exception to the minimal vesicular trafficking machinery of *Giardia*"

### Supplementary Figure 1

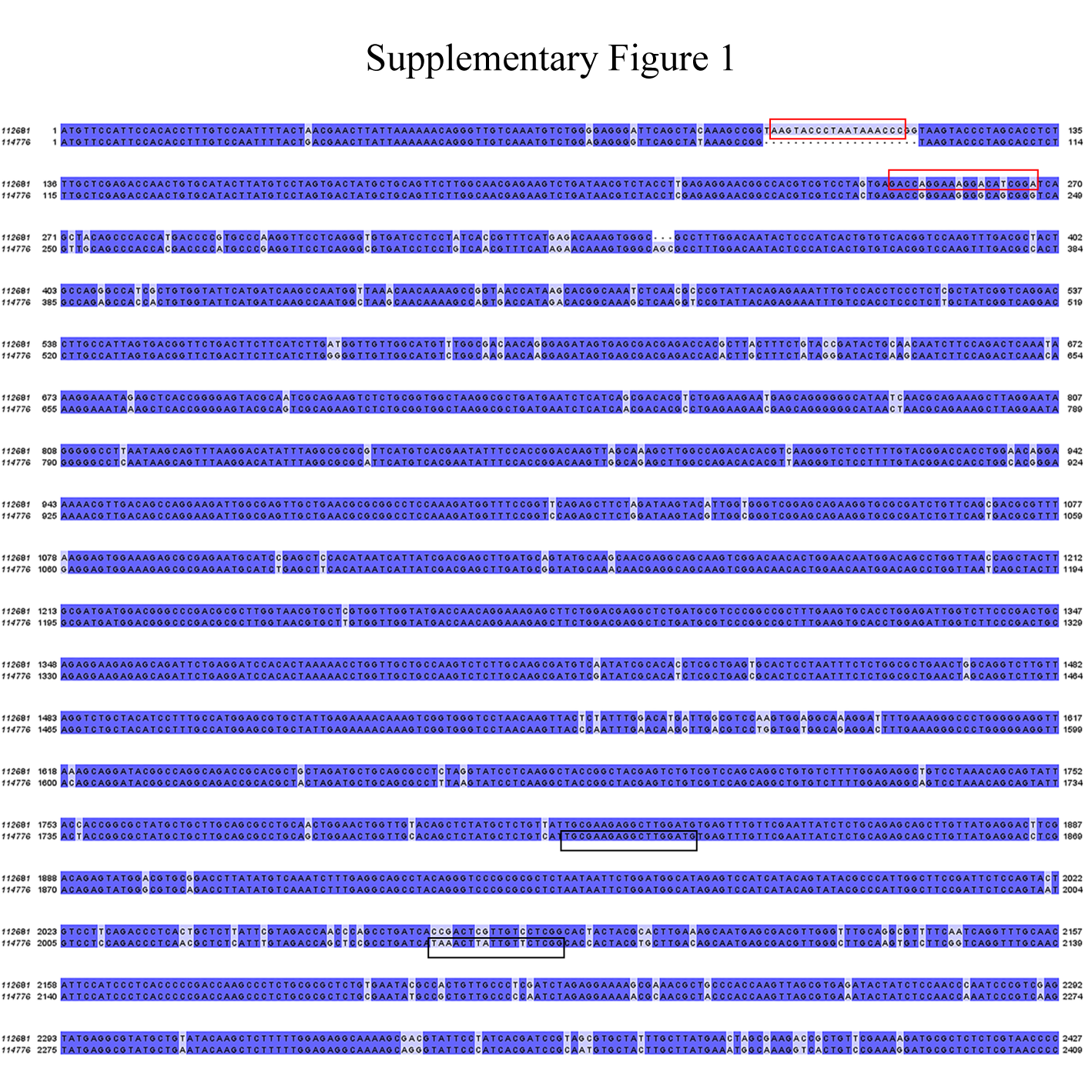

### Supplementary Figure 2

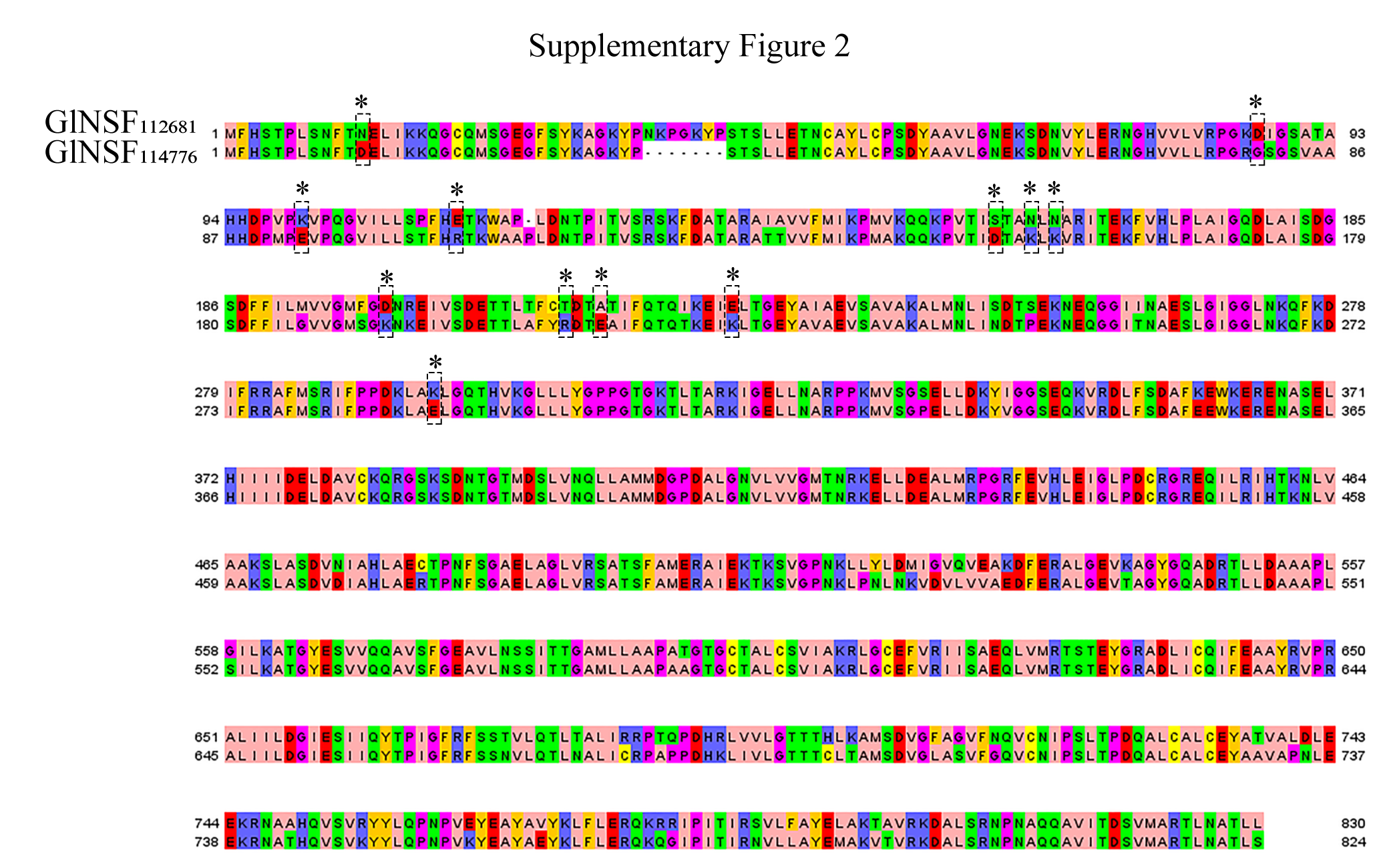

### Supplementary Figure 3

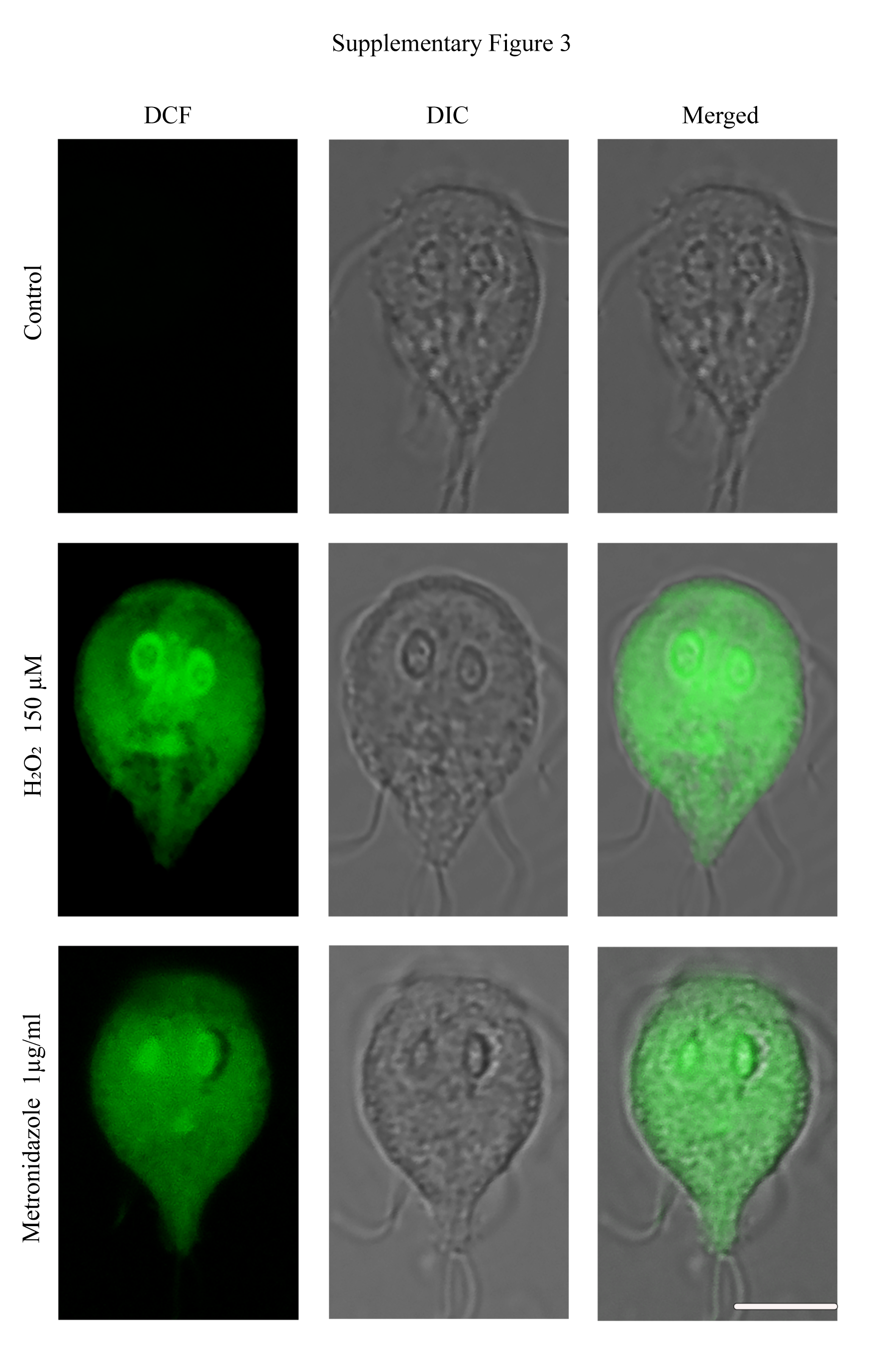

### Supplementary Figure 4

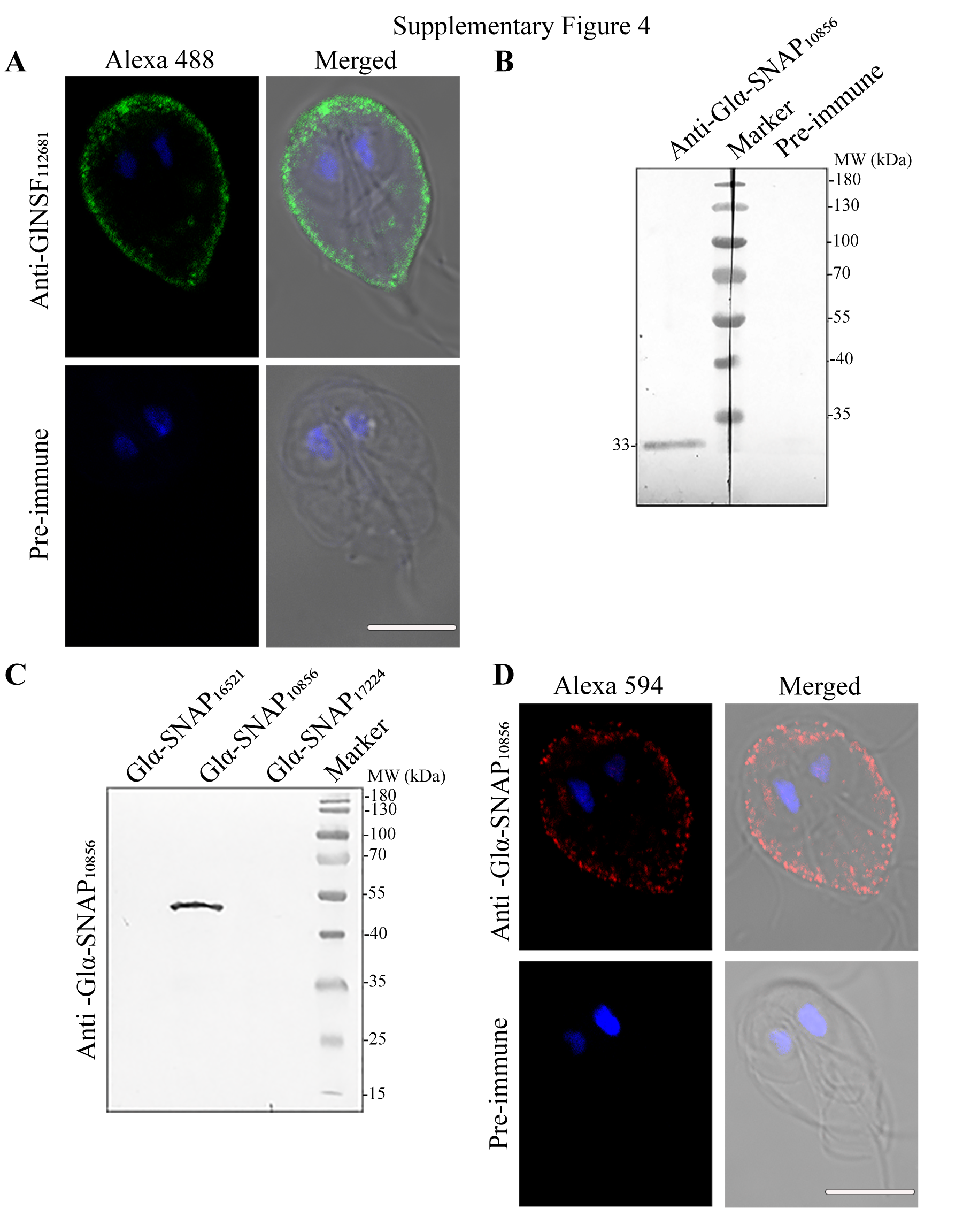
