## Supplementary Table 1 for "Two paralogues of N-ethylmaleimide sensitive factor: An exception to the minimal vesicular trafficking machinery of *Giardia*"

**Supplementary Table 1: List of Primers used in the study**

| Sl No | Primer Sequence (5’ to 3’) | Purpose |
| --- | --- | --- |
| 1 | CAGAATTCGCACTTCATCCGTGTCAGATGC | Forward primer for cloning Glα-SNAP_10856_ in pET32a |
| 2 | GACAAGCTTTCACTGCTCAAAGGTACTTCTCGGC | Reverse primer for cloning Glα-SNAP_10856_ in pET32a |
| 3 | CGGGATCCATGTTCCATTCC | Forward primer for cloning GlNSF_112681_ in pET32a |
| 4 | ACGAATTCTCCAGGTGGTCC | Reverse primer for cloning GlNSF_112681_ in pET32a |
| 5 | CGTGGAATTCGGCTTCAGGCAGATGTCTGACTACG | Forward primer for cloning Glα-SNAP_17224_ in pGBT9 |
| 6 | GCTAGGATCCGCTAAGTCGGCACTGATGTGACATTG | Reverse primer for cloning Glα-SNAP_17224_ in pGBT9 |
| 7 | GAGGGGATCCACAACATGAGTTATGCAAAGCAGGC  GG | Forward primer for cloning Glα-SNAP_16521_ in pGBT9 |
| 8 | CCTTGTCGACCGTTGACGTTGTCCTCGTTCATC | Reverse primer for cloning Glα-SNAP_16521_ in pGBT9 |
| 9 | CAGAATTCGCACTTCATCCGTGTCAGATGC | Forward primer for cloning Glα-SNAP_10856_ in pGBT9 |
| 10 | GACGGATCCTCACTGCTCAAAGGTACTTCTCG | Reverse primer for cloning Glα-SNAP_10856_ in pGBT9 |
| 11 | CGGAGAATTCATGTTCCATTCCACACCTTTGTCC | Forward primer  for cloning GlNSF_114776_ in pGAD424 and pGBT9 |
| 12 | AGTAGTCGACGTTACCAGTTCAGCAGACTCTTGTG | Reverse primer  for cloning GlNSF_114776_ in pGAD424 and pGBT9 |
| 13 | CGGAGAATTCATGTTCCATTCCACACCTTTGTCC | Forward primer  for cloning GlNSF_112681_ in pGAD424 and pGBT9 |
| 14 | AGTAGTCGACGTTACCAGTTCAGCAGACTCTTGTG | Reverse primer  for cloning GlNSF_112681_ in pGAD424 and pGBT9 |
| 15 | AAGTACCCTAATAAACCC | Forward primer  for RT-PCR of GlNSF112681 |
| 16 | ATCCGATGTCCTTTCCTGGTC | Reverse primer  for RT-PCR of  GlNSF112681 |
| 18 | TGCGAAGAGGCTTGGATG | Forward primer  for RT-PCR of  GlNSF114776 |
| 19 | CCGCCAACCCTAAGTTTA | Reverse primer  for RT-PCR of  GlNSF114776 |

**All restriction sites and mutated bases are underlined.**
