## Supplementary Table 2 for "Two paralogues of N-ethylmaleimide sensitive factor: An exception to the minimal vesicular trafficking machinery of *Giardia*"

**Supplementary Table 2: List of peptide fragments that uniquely aligned to GlNSF_114776_**

| Sl No. | Peptide Sequence | Mass | Charge | Abundance |
| --- | --- | --- | --- | --- |
| 1 | AGKYPSTSLLETNCAYLCPSDYAAVLGNEK | 3219.5469 | 5 | 704.77 |
| 2 | ALGEVTAGYGQADR | 1658.8988 | 2 | 3.19e+004 |
| 3 | ALGEVTAGYGQADRTLLDAAAPLSILK | 3156.5819 | 3 | 3768.52 |
| 4 | ALMNLINDTPEK | 1371.7260 | 2 | 1.03e+005 |
| 5 | ALMNLINDTPEKNEQGGITNAESLGIGGLNK | 3580.7271 | 4 | 1.60e+004 |
| 6 | ATTVVFMIKPMAK | 1557.7864 | 3 | 2.09e+004 |
| 7 | TTVVFMIKPMAKQQKPVTIDTAK | 2885.4253 | 4 | 1.36e+004 |
| 8 | DALSRNPNAQQAVITDSVMAR | 2539.2208 | 3 | 2.14e+004 |
| 9 | DLFSDAFEEWK | 1427.5855 | 2 | 2.55e+004 |
| 10 | DLFSDAFEEWKER | 1768.7360 | 3 | 5.19e+004 |
| 11 | DTEAIFQTQTKEIK | 1976.0770 | 3 | 5.52e+004 |
| 12 | DTEAIFQTQTK | 1570.8261 | 3 | 8.36e+004 |
| 13 | EIKLTGEYAVAEVSAVAK | 2013.0108 | 4 | 1017.31 |
| 14 | EIVSDETTLAFYR | 1782.9024 | 3 | 7859.96 |
| 15 | EIVSDETTLAFYRDTEAIFQTQTK | 3172.4846 | 5 | 5.09e+004 |
| 16 | EIVSDETTLAFYRDTEAIFQTQTKEIK | 3273.6414 | 4 | 2388.29 |
| 17 | FDATARATTVVFMIKPMAK | 2155.1295 | 4 | 1.08e+004 |
| 18 | FHSTPLSNFTDELIK | 2038.0736 | 3 | 6.38e+004 |
| 19 | FSSNVLQTLNALICRPAPPDHK | 2524.1968 | 4 | 1.68e+004 |
| 20 | FVHLPLAIGQDLAISDGSDFFILGVVGMSGKNK | 3618.7602 | 5 | 232.95 |
| 21 | GSGSVAAHHDPMPEVPQGVILLSTFHR | 3201.5985 | 4 | 3.04e+004 |
| 22 | GSGSVAAHHDPMPEVPQGVILLSTFHRTK | 3307.4920 | 3 | 3835.58 |
| 23 | IFPPDKLAELGQTHVK | 1848.0035 | 2 | 7.43e+004 |
| 24 | LAELGQTHVK | 1377.8174 | 3 | 1.14e+004 |
| 25 | LAELGQTHVKGLLLYGPPGTGK | 2346.2110 | 3 | 1.21e+004 |
| 26 | LPNLNKVDVLVVAEDFER | 2111.0944 | 4 | 1.05e+005 |
| 27 | LTGEYAVAEVSAVAK | 1869.9951 | 2 | 9.19e+004 |
| 28 | LTGEYAVAEVSAVAKALMNLINDTPEK | \| 2862.5344 | 3 | 6408.75 |
| 29 | MFHSTPLSNFTDELIK | 1920.9364 | 3 | 8.30e+004 |
| 30 | MFHSTPLSNFTDELIKK | 2339.1532 | 4 | 1.83e+005 |
| 31 | MVSGPELLDK | 1101.5592 | 2 | 1.63e+004 |
| 32 | MVSGPELLDKYVGGSEQK | 2188.1880 | 3 | 1.70e+004 |
| 33 | NATHQVSVK | 1265.6602 | 2 | 2.18e+004 |
| 34 | NATHQVSVKYYLQPNPVK | 2207.0409 | 3 | 4.86e+004 |
| 35 | NEQGGITNAESLGIGGLNK | 1884.9893 | 4 | 9708.39 |
| 36 | NEQGGITNAESLGIGGLNKQFK | 2564.3052 | 4 | 1.42e+004 |
| 37 | NKEIVSDETTLAFYR | 1841.0050 | 3 | 3.47e+004 |
| 38 | NKEIVSDETTLAFYRDTEAIFQTQTK | 3141.5113 | 3 | 2718.93 |
| 39 | NLVAAKSLASDVDIAHLAER | 2382.3050 | 4 | 2.00e+005 |
| 40 | NPNAQQAVITDSVMAR | 1966.0634 | 3 | 1.58e+004 |
| 41 | NVLLAYEMAK | 1360.8202 | 3 | 3.31e+004 |
| 42 | NPNAQQAVITDSVMARTLNATLS | 2536.2167 | 3 | 3.36e+004 |
| 43 | NVLLAYEMAKVTVR | 1727.9055 | 2 | 1.36e+004 |
| 44 | QGIPITIR | 910.5450 | 2 | 2.35e+004 |
| 45 | QGIPITIRNVLLAYEMAK | 2231.1358 | 2 | 2985.05 |
| 46 | QKQGIPITIR | 1152.7134 | 2 | 1.76e+005 |
| 47 | QQKPVTIDTAK | 1283.7117 | 2 | 1.37e+004 |
| 48 | QQKPVTIDTAKLK | 1642.8462 | 4 | 2.01e+004 |
| 49 | RNATHQVSVK | 1260.6025 | 2 | 2.27e+004 |
| 50 | SDNVYLERNGHVVLLRPGR | 2263.1942 | 3 | 2.74e+004 |
| 51 | SLASDVDIAHLAER | 1551.7580 | 3 | 3.41e+004 |
| 52 | SLASDVDIAHLAERTPNFSGAELAGLVR | 3002.4867 | 4 | 2.21e+004 |
| 53 | SVGPNKLPNLNK | 1415.7728 | 2 | 4969.71 |
| 54 | TKWAAPLDNTPITVSR | 1862.9402 | 3 | 1.14e+004 |
| 55 | TLLDAAAPLSILK | 1460.7539 | 2 | 1087.42 |
| 56 | TLNATLL | 1017.5875 | 2 | 1.77e+004 |
| 57 | VDVLVVAEDFER | 1403.7469 | 2 | 2.04e+004 |
| 58 | VRDLFSDAFEEWK | 1762.7573 | 2 | 1988.59 |
| 59 | VRDLFSDAFEEWKER | 2220.1275 | 4 | 7.70e+004 |
| 60 | WAAPLDNTPITVSR | 1633.7853 | 3 | 1.37e+004 |
| 61 | WAAPLDNTPITVSRSK | 1768.8405 | 3 | 3.09e+004 |
| 62 | YEAYAEYK | 1077.4783 | 2 | 2985.17 |
| 63 | YVGGSEQK | 1118.6563 | 2 | 1.92e+005 |
| 64 | YVGGSEQKVR | 1201.6190 | 3 | 2.79e+004 |
| 65 | YYLQPNPVK | 1410.7474 | 3 | 7670.32 |
| 66 | YYLQPNPVKYEAYAEYK | 2302.0603 | 2 | 3.87e+004 |

**Sequence coverage of GlNSF_114776_ by unique aligned reads**

MFHSTPLSNFTDELIKKQGCQMSGEGFSYKAGKYPSTSLLETNCAYLCPSDYAAVLGNEKSDNVYLERNGHVVLLRPGRGSGSVAAHHDPMPEVPQGVILLSTFHRTKWAAPLDNTPITVSRSKFDATARATTVVFMIKPMAKQQKPVTIDTAKLKVRITEKFVHLPLAIGQDLAISDGSDFFILGVVGMSGKNKEIVSDETTLAFYRDTEAIFQTQTKEIKLTGEYAVAEVSAVAKALMNLINDTPEKNEQGGITNAESLGIGGLNKQFKDIFRRAFMSRIFPPDKLAELGQTHVKGLLLYGPPGTGKTLTARKIGELLNARPPKMVSGPELLDKYVGGSEQKVRDLFSDAFEEWKERENASELHIIIIDELDAVCKQRGSKSDNTGTMDSLVNQLLAMMDGPDALGNVLVVGMTNRKELLDEALMRPGRFEVHLEIGLPDCRGREQILRIHTKNLVAAKSLASDVDIAHLAERTPNFSGAELAGLVRSATSFAMERAIEKTKSVGPNKLPNLNKVDVLVVAEDFERALGEVTAGYGQADRTLLDAAAPLSILKATGYESVVQQAVSFGEAVLNSSITTGAMLLAAPAAGTGCTALCSVIAKRLGCEFVRIISAEQLVMRTSTEYGRADLICQIFEAAYRVPRALIILDGIESIIQYTPIGFRFSSNVLQTLNALICRPAPPDHKLIVLGTTTCLTAMSDVGLASVFGQVCNIPSLTPDQALCALCEYAAVAPNLEEKRNATHQVSVKYYLQPNPVKYEAYAEYKLFLERQKQGIPITIRNVLLAYEMAKVTVRKDALSRNPNAQQAVITDSVMARTLNATLS
