## Supplementary Table 3 for "Two paralogues of N-ethylmaleimide sensitive factor: An exception to the minimal vesicular trafficking machinery of *Giardia*"

**Supplementary Table 3: List of peptide fragments that uniquely aligned to GlNSF_112681_**

| Sl No. | Peptide Sequence | Mass | Charge | Abundance |
| --- | --- | --- | --- | --- |
| 1 | AGKYPNKPGK | 1268.5900 | 2 | 1.15e+005 |
| 2 | AGYGQADRTLLDAAAPLGILK | 2277.1580 | 3 | 1.84e+005 |
| 3 | AIAVVFMIKPMVK | 1545.8683 | 3 | 4402.03 |
| 4 | AIAVVFMIKPMVKQQKPVTISTANLNAR | 3319.8323 | 6 | 1845.54 |
| 5 | ALGEVK | 657.3687 | 2 | 6856.59 |
| 6 | ALGEVKAGYGQADR | 1489.8006 | 3 | 8.15e+004 |
| 7 | ALMNLISDTSEK | 1334.6886 | 2 | 8.43e+004 |
| 9 | ATGYESVVQ | 959.5660 | 2 | 1855.32 |
| 10 | DFERALGEVK | 1414.7054 | 3 | 3.98e+004 |
| 11 | DIGSATAHHDPVPK | 1457.7280 | 2 | 5.37e+004 |
| 12 | DLFSDAFKEWK | 1384.7073 | 3 | 2.86e+004 |
| 13 | DLFSDAFKEWKER | 1959.9555 | 2 | 1.88e+004 |
| 14 | EIELTGEYAIAEVSAVAK | 2262.0845 | 3 | 1.81e+004 |
| 15 | EIELTGEYAIAEVSAVAKALMNLISDTSEK | 3294.6355 | 4 | 74.32 |
| 16 | EIVSDETTLTFCTDTATIFQTQIK | 2846.3862 | 3 | 1.83e+004 |
| 17 | FDATARAIAVVFMIKPMVK | 2229.1675 | 3 | 8880.91 |
| 18 | FHSTPLSNFTNELIK | 2110.1046 | 4 | 2.56e+004 |
| 19 | FSSTVLQTLTALIR | 1684.6849 | 2 | 1914.45 |
| 20 | FSSTVLQTLTALIRRPTQPDHR | 2536.2023 | 3 | 1.21e+004 |
| 21 | FVHLPLAIGQDLAISDGSDFFILMVVGMFGDNR | 3956.9724 | 5 | 1.74e+004 |
| 22 | IFPPDKLAK | 1027.5335 | 2 | 1.64e+004 |
| 23 | IGELLNARPPKMVSGSELLDK | 2404.1971 | 3 | 1.03e+005 |
| 24 | IPITIRSVLFAYELAK | 1955.0008 | 5 | 3.99e+005 |
| 25 | KQGCQMSGEGFSYK | 1562.6862 | 3 | 2.00e+004 |
| 26 | LAKLGQTHVK | 1135.7082 | 1 | 4.36e+004 |
| 27 | LLYLDMIGVQVEAK | 1646.8960 | 3 | 3.58e+004 |
| 28 | LLYLDMIGVQVEAKDFER | 2232.0598 | 5 | 5.69e+004 |
| 29 | LVVLGTTTHLK | 1264.7173 | 2 | 4.48e+004 |
| 30 | MFHSTPLSNFTNELIK | 1971.9236 | 3 | 2.97e+004 |
| 31 | MFHSTPLSNFTNELIKK | 2105.9839 | 4 | 3.23e+005 |
| 32 | MVSGSELLDK | 1077.5301 | 2 | 1.10e+005 |
| 33 | MVSGSELLDKYIGGSEQK | 2150.1455 | 4 | 1.31e+004 |
| 34 | NAAHQVSVR | 994.5257 | 2 | 9444.57 |
| 35 | NAAHQVSVRYYLQPNPVEYEAYAVYK | 3434.7148 | 4 | 5.45e+005 |
| 36 | NEQGGIINAESLGIGGLNK | 1896.9499 | 4 | 1.95e+004 |
| 37 | NEQGGIINAESLGIGGLNKQFK | 2380.2103 | 4 | 2.07e+004 |
| 38 | NGHVVLVRPGKDIGSATAHHDPVPK | 2684.3990 | 3 | 4.28e+004 |
| 39 | QQKPVTISTANLNAR | 1719.8615 | 3 | 1.12e+004 |
| 40 | QQKPVTISTANLNARITEK | 2271.1238 | 3 | 1.20e+004 |
| 41 | RNAAHQVSVR | 1258.4932 | 2 | 6.43e+004 |
| 42 | RPTQPDHR | 1099.5208 | 2 | 2.27e+004 |
| 43 | RPTQPDHRLVVLGTTTHLK | 2262.2325 | 3 | 2.56e+004 |
| 44 | SDNVYLERNGHVVLVRPGK | 2249.0451 | 5 | 6.62e+004 |
| 45 | SLASDVNIAHLAECTPNFSGAELAGLVR | 3106.6505 | 3 | 2.19e+004 |
| 46 | SVGPNKLLYLDMIGVQVEAK | 2295.1443 | 4 | 2.13e+004 |
| 47 | SVLFAYELAK | 1181.6064 | 2 | 1.28e+005 |
| 48 | SVLFAYELAKTAVR | 1702.8595 | 3 | 8.79e+004 |
| 49 | TLLDAAAPLGILK | 1504.7083 | 3 | 1241.56 |
| 50 | VPQGVILLSPFHETK | 1757.8583 | 4 | 7.10e+004 |
| 51 | VRDLFSDAFK | 1318.5731 | 3 | 8159.19 |
| 52 | VRDLFSDAFKEWK | 1653.8462 | 3 | 1.25e+004 |
| 53 | VRDLFSDAFKEWKER | 1924.9215 | 3 | 2.92e+004 |
| 54 | VRDLFSDAFKEWKERENASELHIIIIDELDAVCK | 4195.0484 | 5 | 1.03e+004 |
| 55 | WAPLDNTPITVSR | 1633.7853 | 3 | 1.37e+004 |
| 56 | WAPLDNTPITVSRSK | 1777.8633 | 3 | 5.13e+004 |
| 57 | YIGGSEQK | 1285.7316 | 2 | 1.96e+004 |
| 58 | YIGGSEQKVR | 1135.5898 | 2 | 5.68e+004 |
| 59 | YPNKPGKYPSTSLLETNCAYLCPSDYAAVLGNEK | 3827.7053 | 5 | 9840.29 |
| 60 | YYLQPNPVEYEAYAVYK | 2109.0003 | 3 | 5.79e+004 |
| 61 | YYLQPNPVEYEAYAVYKLFLER | 2781.3467 | 3 | 9.68e+004 |

**Sequence coverage of GlNSF_112681_ by unique aligned reads**

MFHSTPLSNFTNELIKKQGCQMSGEGFSYKAGKYPNKPGKYPSTSLLETNCAYLCPSDYAAVLGNEKSDNVYLERNGHVVLVRPGKDIGSATAHHDPVPKVPQGVILLSPFHETKWAPLDNTPITVSRSKFDATARAIAVVFMIKPMVKQQKPVTISTANLNARITEKFVHLPLAIGQDLAISDGSDFFILMVVGMFGDNREIVSDETTLTFCTDTATIFQTQIKEIELTGEYAIAEVSAVAKALMNLISDTSEKNEQGGIINAESLGIGGLNKQFKDIFRRAFMSRIFPPDKLAKLGQTHVKGLLLYGPPGTGKTLTARKIGELLNARPPKMVSGSELLDKYIGGSEQKVRDLFSDAFKEWKERENASELHIIIIDELDAVCKQRGSKSDNTGTMDSLVNQLLAMMDGPDALGNVLVVGMTNRKELLDEALMRPGRFEVHLEIGLPDCRGREQILRIHTKNLVAAKSLASDVNIAHLAECTPNFSGAELAGLVRSATSFAMERAIEKTKSVGPNKLLYLDMIGVQVEAKDFERALGEVKAGYGQADRTLLDAAAPLGILKATGYESVVQQAVSFGEAVLNSSITTGAMLLAAPATGTGCTALCSVIAKRLGCEFVRIISAEQLVMRTSTEYGRADLICQIFEAAYRVPRALIILDGIESIIQYTPIGFRFSSTVLQTLTALIRRPTQPDHRLVVLGTTTHLKAMSDVGFAGVFNQVCNIPSLTPDQALCALCEYATVALDLEEKRNAAHQVSVRYYLQPNPVEYEAYAVYKLFLERQKRRIPITIRSVLFAYELAKTAVRKDALSRNPNAQQ
