## Supplementary Table 4 for "Two paralogues of N-ethylmaleimide sensitive factor: An exception to the minimal vesicular trafficking machinery of *Giardia*"

**Supplementary Table 4 : List of constructs used in the study**

| Construct name | Cloned gene | Description | Primers used |
| --- | --- | --- | --- |
| pTG1 | GlNSF_112681_ | N-terminal of GlNSF_112681_ (pET32a) | 3 and 4 |
| pTG2 | Glα-SNAP_10856_ | Glα-SNAP_10856_ (pET32a) | 1 and 2 |
| pTG3 | Glα-SNAP_10856_ | Glα-SNAP_10856_  (pGBT9) | 9 and 10 |
| pTG4 | Glα-SNAP_16521_ | Glα-SNAP_16521_ (pGBT9) | 7 and 8 |
| pTG5 | Glα-SNAP_17224_ | Glα-SNAP_17224_ (pGBT9) | 5 and 6 |
| pTG6 | GlNSF_112681_ | GlNSF_112681_  (pGBT9) | 13 and 14 |
| pTG7 | GlNSF_112681_ | GlNSF_112681_  (pGAD424) | 13 and 14 |
| pTG8 | GlNSF_114776_ | GlNSF_114776_  (pGBT9) | 11 and 12 |
| pTG9 | GlNSF_114776_ | GlNSF_114776_  (pGAD424) | 11 and 12 |
| pTG10 | Recombinant  GlNSF_114776_ | GlNSF_114776_ (pGAD424) – N terminal domain of GlNSF_112681_ and remainder GlNSF_114776_ | Created using common restriction sites Bsu36I and EcoRI in both pTG9 and pTG7 construct |
| pTG11 | Recombinant  GlNSF_112681_ | GlNSF_112681_ (pGAD424) – N terminal domain of GlNSF_114776_ and remainder GlNSF_112681_ | Created using common restriction sites Bsu36I and EcoRI in both pTG9 and pTG7 construct |
